# PIPI: PTM-Invariant Peptide Identification Using Coding Method

**DOI:** 10.1101/055806

**Authors:** Fengchao Yu, Ning Li, Weichuan Yu

## Abstract

In computational proteomics, identification of peptides with an unlimited number of post-translational modification (PTM) types is a challenging task. The computational cost increases exponentially with respect to the number of modifiable amino acids and linearly with respect to the number of potential PTM types at each amino acid. The problem becomes intractable very quickly if we want to enumerate all possible modification patterns. Existing tools (e.g., MS-Alignment, ProteinProspector, and MODa) avoid enumerating modification patterns in database search by using an alignment-based approach to localize and characterize modified amino acids. This approach avoids enumerating all possible modification patterns in a database search. However, due to the large search space and PTM localization issue, the sensitivity of these tools is low. This paper proposes a novel method named PIPI to achieve PTM-invariant peptide identification. PIPI first codes peptide sequences into Boolean vectors and converts experimental spectra into real-valued vectors. Then, it finds the top 10 peptide-coded vectors for each spectrum-coded vector. After that, PIPI uses a dynamic programming algorithm to localize and characterize modified amino acids. Simulations and real data experiments have shown that PIPI outperforms existing tools by identifying more peptide-spectrum matches (PSMs) and reporting fewer false positives. It also runs much faster than existing tools when the database is large.

## 1 Introduction

Shotgun proteomics has achieved great success after more than 20 years’ development. Based on the database search idea, researchers have proposed many tools to identify peptides. According to the approaches to dealing with post-translational modification (PTM), we can classify these tools into two categories: restricted tools^1–15^ and unrestricted tools^5,16–35^.

Restricted tools generate theoretical spectra by *in silico* fragmenting peptide sequences. They infer an experimental spectrum’s corresponding peptide sequence by finding the most similar theoretical spectrum. These tools need to generate different theoretical spectra corresponding to different modification patterns. Given a peptide sequence, the number of theoretical spectra follows

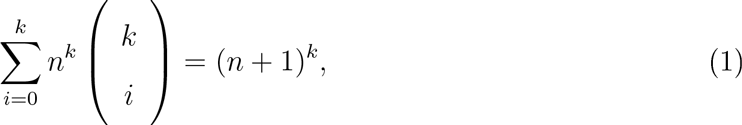
where *k* is the number of modiable amino acids and *n* is the average number of potential PTM types at each modifiable amino acid. The number of theoretical spectra already becomes very large, even when we only consider a few PTM types. Some tools^9–15^use tags to accelerate searching speed. A tag is a sequence fragment inferred from a spectrum, and based on the number of amino acids, they can have various lengths. For simplicity, from now on, we use “spectrum” to refer to “experimental spectrum” if there is no possible confusion. Given a spectrum, tag-based tools infer the tag compositions and locate peptide sequences containing those tags, and they use these peptide sequences as a custom database to search for the result. Even with optimized algorithms, the problem becomes intractable very quickly if we want to enumerate all modification patterns. Thus, these tools only allow a small number of modified amino acids and PTM types during a database search.

Unrestricted tools identify spectra with unlimited PTM types by inferring the locations of modified amino acids during a database search. Spider^18^ and OpenSea^20^ obtain parts of a spectrum’s sequence by *de novo* sequencing^36–38^. Then, they infer the modified amino acids by comparing the sequence parts with the corresponding peptide sequence from a database. MS-Alignment^17^ compares an experimental spectrum with PTM-free theoretical spectra. It uses a dynamic programming algorithm with five jumping rules to find the overlapping peaks, and treats gaps between the overlapping parts as modified amino acids. MS-Alignment only supports up to two modifiable amino acids in each spectrum. MODa^31^ infers various lengths of sequence fragments from a spectrum, and aligns tags against peptide sequences. It also uses a dynamic programming algorithm to find the best alignment result. After the alignment, it calculates a score for each sequence and selects the one with the highest score.

All these tools’ scoring functions rely on the modification pattern, which means that the accuracy of PTM localization strongly influences the performance of the identification. However, PTM localization is not an easy task. Although various methods have been proposed^39–41^, it is still difficult to determine the exact locations^42–44^.

In this paper, we propose a PTM-invariant peptide identification method named PIPI. which belongs to the category of unrestricted tools. PI PI first builds a theoretical database of peptide sequences by converting each sequence into a coded Boolean vector. Each element in the vector indicates whether the corresponding three-amino-acid tag exists in the original sequence. When analyzing a spectrum, PIPI only extracts peaks whose relative distances are invariant to PTM. Then, it converts peaks into a vector and searches for the top 10 candidates. PIPI doesn’t need to infer the exact locations of modified amino acids during the database search. Thus, it bypasses the PTM localization problem in database search and leads to a better performance in both peptide identification and PTM characterization.

**Figure 1:**
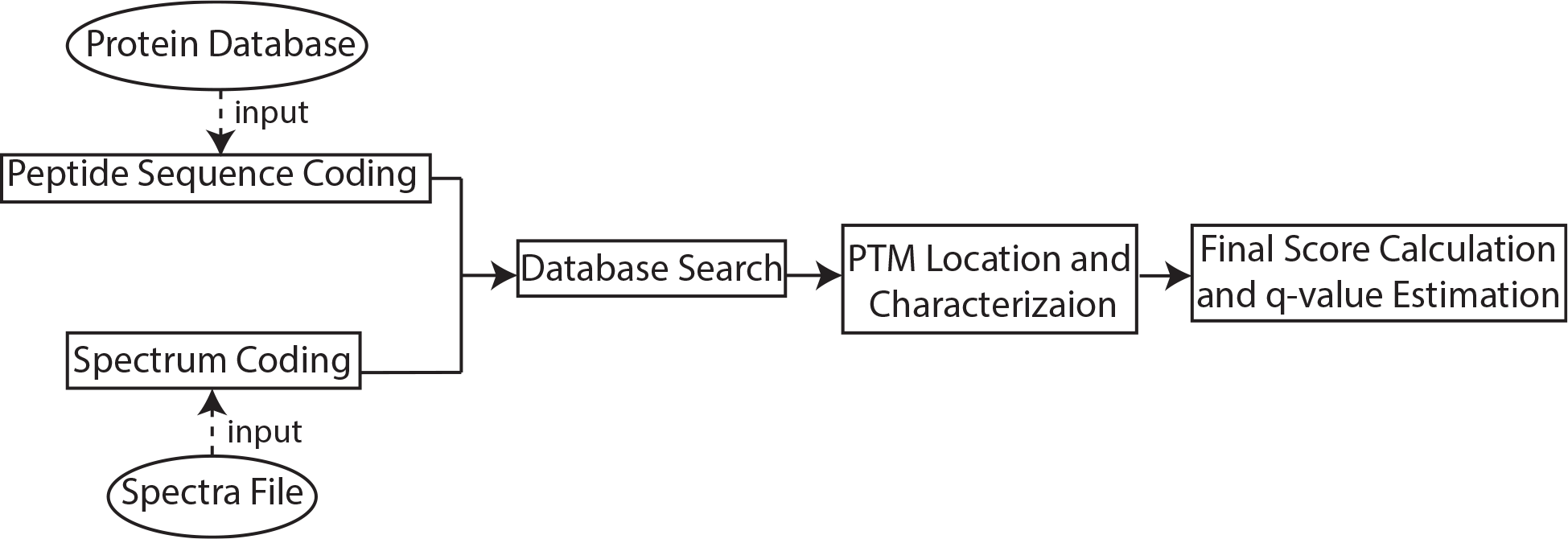
The work-flow of PIPI.

The rest of the paper is organized as follows: Section 2 describes coding, database search. PTM localization and characterization, final score calculation, and *q*-value estimation in detail. Section 3 presents three sets of experiments to demonstrate the performance of PIPI. Section 4 discusses the relationship between PIPI and existing tools. It also raises the issue of low accuracy in PTM localization and characterization.

## 2 Methodology

Figure 1 shows the work-flow of PIPI. There are four major steps:

1. Peptide sequence coding and spectrum coding.
2. Database search.
3. PTM location and characterization.
4. Final score calculation and g-value estimation.

We will describe each step in detail.

**Figure 2:**
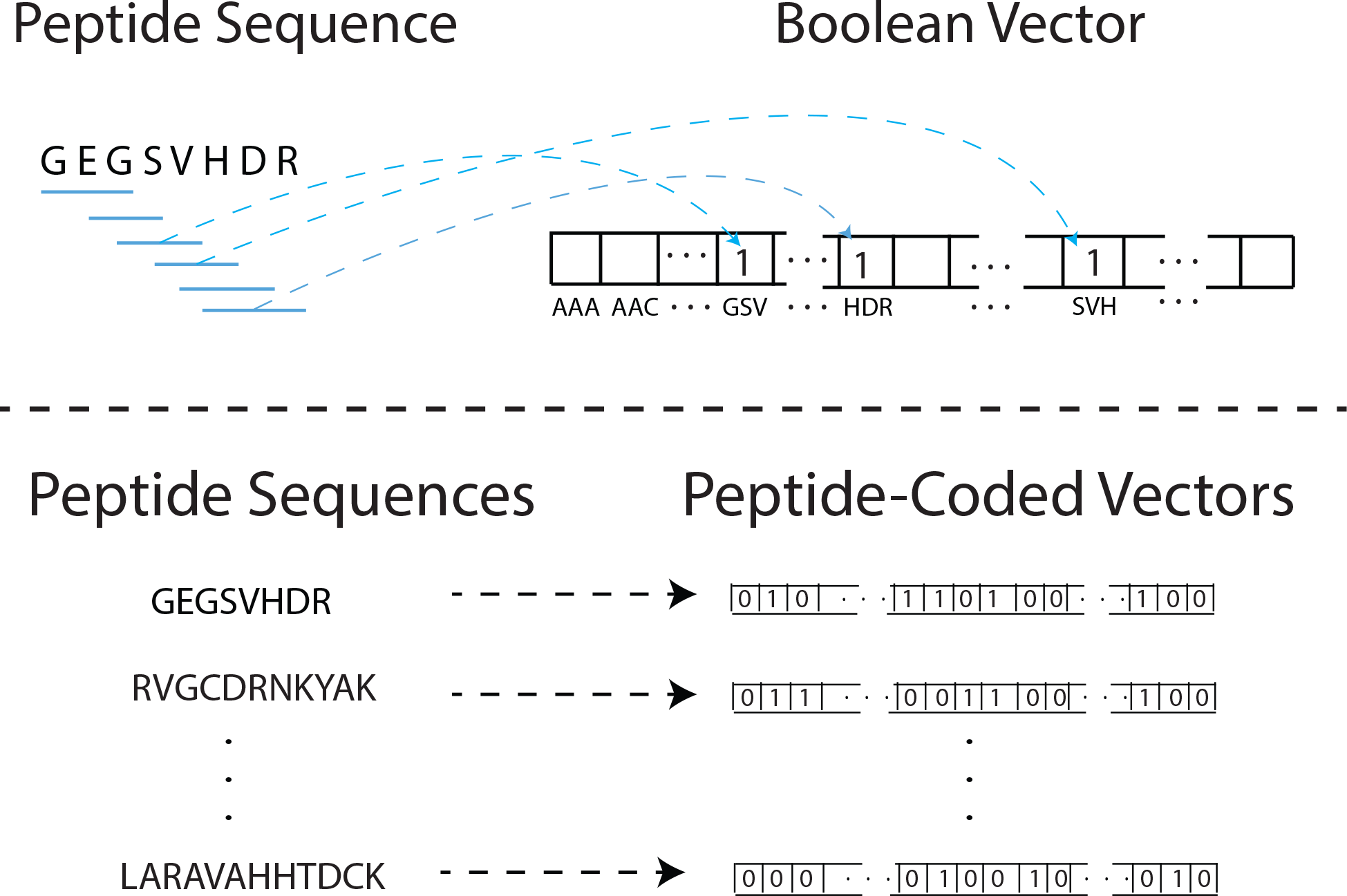
An illustration of peptide coding and database building. **Top:** PIPI extracts tags from a peptide sequence with an overlapped sliding window. It codes different tags into a Boolean vector. Each peptide sequence in the database becomes a coded vector. **Bottom:** PIPI codes all peptide sequences into Boolean vectors one by one.

## 2.1 Peptide sequence coding and spectrum coding

### 2.1.1 Peptide sequence coding

PIPI *in silico* digests proteins to peptides and codes peptide sequences into Boolean vectors, called peptide-coded vectors. Given a peptide sequence. PIPI extracts three-length tags from the N-terminal to the C-terminal with an overlapped sliding window (Figure 2). Because there is 110 intensity information in a peptide sequence, values in the peptide-coded vector are either 0 or 1. Elements corresponding to extracted tags have 1 values and the others have 0 values. PIPI codes each digested peptide sequence into a peptide-coded vector, and after coding, it indexes all vectors based 011 their corresponding peptides’ masses.

### 2.1.2 Spectrum coding

Given a spectrum, PIPI first removes noisy peaks and normalizes peak intensities. It uses the peak intensity with the highest frequency as a threshold^45^, and eliminates all peaks whose intensities are smaller than the threshold. Then, PIPI replaces each peak’s intensity with its square root and normalizes the peak intensities, as in SEQUEST^2^, by dividing the whole *m*/*z* range into 10 regions. In each region, it normalizes peak intensities so that the highest one equals 1.

Some ions may not be detected in a spectrum, due to the limited fragmentation efficiency and instrument’s detection sensitivity. PIPI checks each peak to see if its complementary peak exists in a spectrum. If not, it adds the complementary peak with the same intensity to the corresponding location. Two peaks are complementary to each other if the sum of their *m*/*z* values equals *S_m_*+2 ×*p_m_*, where *S_m_* is the precursor mass of the spectrum and *p_m_* is the mass of a proton. Please note that PIPI only considers single charged fragmentations. PIPI also adds two one-intensity peaks with the *m*/*z* values corresponding to the N-terminal and the C-terminal, respectively.

After adding peaks, PIPI expresses a spectrum as a matrix **S**_*L_s_*×2_, where *L_s_* is the total number of peaks. The elements ***s***_*i*,1_ and ***s***_*i*,2_ are the *m/z* value and the intensity of the *i*-th peak, respectively. Two peaks can form a peak pair if they satisfy the following relationship |***s***_*j*,1_ − ***s***_*i*,1_ − Δ_*k*_|≤2τ, where *k* ∈ [1, 22] is an index of the 22 amino acids (considering “U” and “O”), Δ_*k*_ is the mass of one of the 22 amino acids, and *τ* is the MS/MS mass tolerance. A peak pair consisting of the *i*-th and the *j*-th peak is denoted as *P*(*i*,*j*). Two peak pairs *P*(*i*,*j*) and *P*(*i*′,*j*′) can be linked if *j*=*i*′, and a number of pairs can be linked sequentially to form a tag. For simplicity, we denote an *L_t_* length tag as *P*_1_*P*_2_… *P*_*L_t_*_. In practice, most spectra can produce many tags due to the large number of noisy peaks. If there were more than 200 tags in a spectrum, PIPI would divide the whole *m*/*z* range into 10 regions and keep the top 20 tags in each region.

PIPI codes tags with the same length into a vector v = [*v*_1_,*v*_2_,…, *v*_*i*_,…, *v*_*L_v_*_], where *L_v_* is the length of the vector and

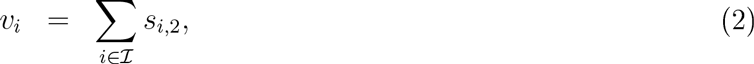

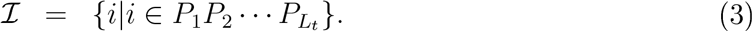

Here *i* ∈ *P*_1_*P*_2_… *P*_*L_t_*_ means that *i* belongs to one of the indexes of the peaks forming *P*_1_*P*_2_… *P*_*L_t_*_. In this paper, we set *L_t_* = 3, and will demonstrate that this is a good choice later on. We call the vector a spectrum-coded vector. Since the tags extracted from a spectrum can be from b-ions or y-ions (under a collision-induced dissociation (CID) setting), PIPI cannot determine the direction of a tag. Thus, PIPI treats a tag and its reversed version as the same. PIPI also treats amino acids “I” and “J” equally because they have an identical mass. There are in total 22 amino acids, including two additional ones, “U” and “O”. With the setting above, we can obtain the length of the vector:

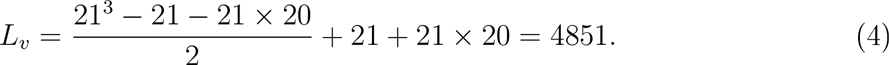

The order of the tags doesn’t matter as long as it is consistent in the whole work-flow. Figure 3 illustrates how PIPI codes a spectrum.

## 2.2 Similarity measure and database search

### 2.2.1 Similarity measure

A spectrum-coded vector contains local information of a spectrum. A peptide-coded vector contains the sequence information of a peptide. PIPI uses the cross-correlation coefficient as the similarity measure:

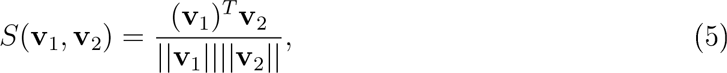
where **v**_1_ is a spectrum-coded vector and **v**_2_ is a peptide-coded vector. There are two parts in the cross-correlation coefficient: a dot product in the numerator and a product of two *l*-2 norms in the denominator. Given two vectors, the dot product measures the overlapping level. The denominator normalizes the dot product, and the cross-correlation coefficient measures the similarity between a spectrum and a peptide sequence.

**Figure 3:**
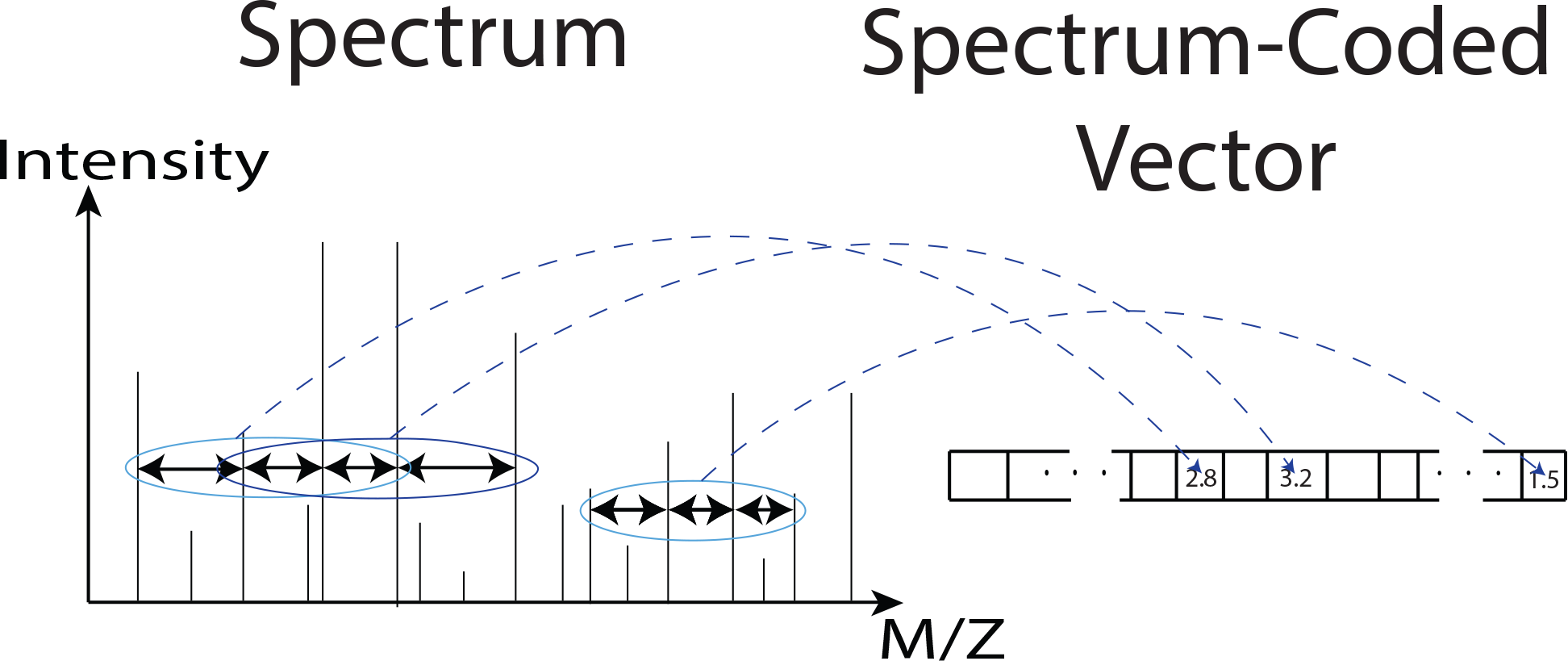
An illustration of spectrum coding. PIPI extracts three-length tags from a spectrum and codes these tags into a vector. Its indexes indicate different tags, and its values are the intensity summations of the peaks forming the corresponding tags.

In order to choose the right tag length, we studied the discriminant power of different lengths empirically. We used the whole proteome of Homo sapiens (human) from UniProtKB/Swiss-Prot (20.205 proteins. 2015–11 release) using *in silico* digested these proteins with trypsin, and kept peptides with masses from −500 Da to 5.000 Da. allowing no missed-cleavage. There were in total 638,480 nonredundant peptides. We let PIPI code all these peptides and calculate the cross-correlation coefficients using pairs of peptide-coded vectors whose peptides’ masses’ differences were from — 250 Da to 250 Da. It used different tag lengths, from 2 to 4 amino acids, for coding. Table 1 shows the relative frequencies of the cross-correlation coefficients from 0 to 0.5. Please note that the cross-correlation coefficient of two identical vectors equals 1. Most of the cross-correlation coefficients under the “tag 3” and “tag 4” settings are smaller than 0.1, which means that PIPI can separate coded vectors from different peptide sequences well. Since a longer tag requires a higher spectrum quality, which is not always satisfied, we decided to use tags of length of three amino acids.

**Table 1.** Relative frequencies of the cross-correlation coefficients from 0 to 0.5.

| Cross-Correlation Coefficients | Tag Lengths | Tag 2 | Tag 3 | Tag 4 |
| --- | --- | --- | --- | --- |
| 0~0.1 |  | 0.6496 | 0.9735 | 0.9982 |
| 0.1~0.2 |  | 0.2168 | 0.0206 | 0.0013 |
| 0.2~0.3 |  | 0.1016 | 0.0050 | 0.0000 |
| 0.3~0.4 |  | 0.0233 | 0.0000 | 0.0000 |
| 0.4~0.5 |  | 0.0067 | 0.0000 | 0.0000 |

We also used a real data set to investigate the discriminant power of coded vectors coupled with the cross-correlation coefficient measure. We chose 18,757 MS/MS spectra from a data set published by Chick et al.^34^. There are 14,843 PTM-free spectra and 3,914 PTM-containing spectra, and all of them were identified by Comet ^7^ with *q*-values≤0.01. Comet is an open source implementation of SEQUEST’s algorithm. We set 5 variable modifications (i.e. Oxidation, Phosphorylation, Acetylation, Methylation, and Deamidated) in using Comet. We let PIPI code them and calculate the cross-correlation coefficients using pairs of coded vectors. Without considering PTM difference, if a pair of two different vectors was from the same peptide, it was called a homologous pair, and if a pair of two different vectors was from different peptides, it was called a heterologous pair. The allowed peptide mass difference was also from −250 Da to 250 Da. Because we allowed a wide mass difference and did not consider PTM difference in determining the homologous pairs and heterologous pairs, the comparison was PTM-invariant. Figure 4 shows two histograms corresponding to crosscorrelation coefficients of the homologous pairs and heterologous pairs, respectively. Each histogram was normalized by its total count. We can see that the coded vectors coupled with the cross-correlation coefficient measure separates the heterologous pairs from homologous pairs well.

**Figure 4:**
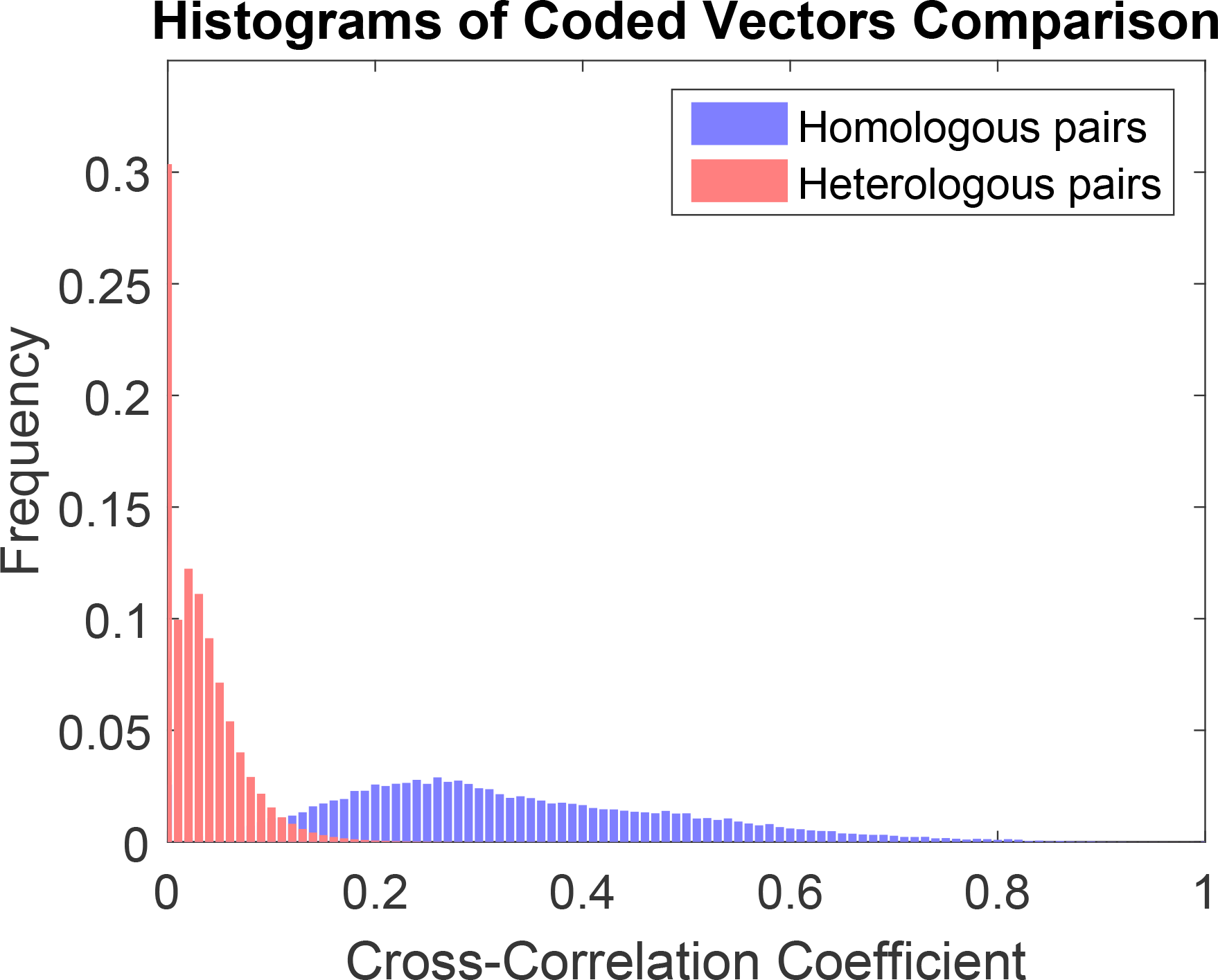
Two histograms of the cross-correlation coefficients from pair-wisely comparing coded vectors. Homologous pairs and heterologous pairs are labeled with different colors.

#### 2.2.2 Database search

After coding all spectra and peptide sequences. PIPI finds the 10 most similar peptide-coded vectors for each spectrum-coded vector. Given a spectrum-coded vector. PIPI first finds all possible peptide-coded vectors whose corresponding peptides’ masses are within the range [*S_m_* − *v*,*S*_m_+*v*], where *S_m_* is the spectrum’s precursor mass and *v* is a pre-defined value. Then, it uses Equation (5) to measure the similarity between a spectrum-coded vector and a peptide-coded vector. For each spectrum-coded vector. PIPI only keeps the top 10 peptide-coded vectors with the highest similarity scores.

### 2.3 PTM location and characterization

Here we describe how PIPI locates and characterizes modified amino acids given a spectrum and a peptide sequence.

Researchers have used dynamic programming based approaches to infer the locations and mass shifts of modified amino acids. MS-Alignment aligns an experimental spectrum’s peaks against those from a theoretical spectrum, while MODa aligns tags of different lengths against a peptide sequence. Since PIPI’s coding procedure has already extracted three-length tags from a spectrum, it aligns tags against a peptide sequence using dynamic programming.

Before the alignment, PIPI compares each tag’s *m*/*z* value in the experimental spectrum with that in the PTM-free theoretical spectrum. It only keeps those tags whose experimental *m*/*z* values are within the range [*T_mz_* − *v*, *T_mz_*+*v*], where *T_mz_* is the *m*/*z* value in the PTM-free theoretical spectrum. We call this process tag cleaning (Figure 5). After tag cleaning, PIPI adds the N-terminal and the C-terminal as two special tags.

We denote a tag as *t_i_* where *i* is an index. We define 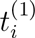 as the location of the first amino acid in the peptide sequence and *I*(*t_i_*) as the summation of the peak intensities of the tag. The dynamic programming matrix is **D**_|{t_i_}|×(*n*+2)_, where |{*t_i_*}| is the number of tags and *n* equals the length of the peptide sequence. During the dynamic programming, there are two kinds of jumps: jumps within a tag and jumps between two tags. Because tags have sequence and peak location information, not all jumps between tags are meaningful. Thus, we define the following jumping rules (Figure 6):

1. Jumps within a tag are allowed (the green arrows in Figure 6).
2. Jumps from the end of a tag to the start of another tag are allowed (the black arrows in Figure 6).
3. Jumps from the middle of a tag to the start of another tag are allowed only if the end of the former tag overlaps with the start of the latter tag (the blue arrow in Figure 6). Overlapping means that they have the same substring and the same peak locations.
4. Jumps from the end of a tag to the end of another tag are not allowed (the red arrow in Figure 6).

**Figure 5:**
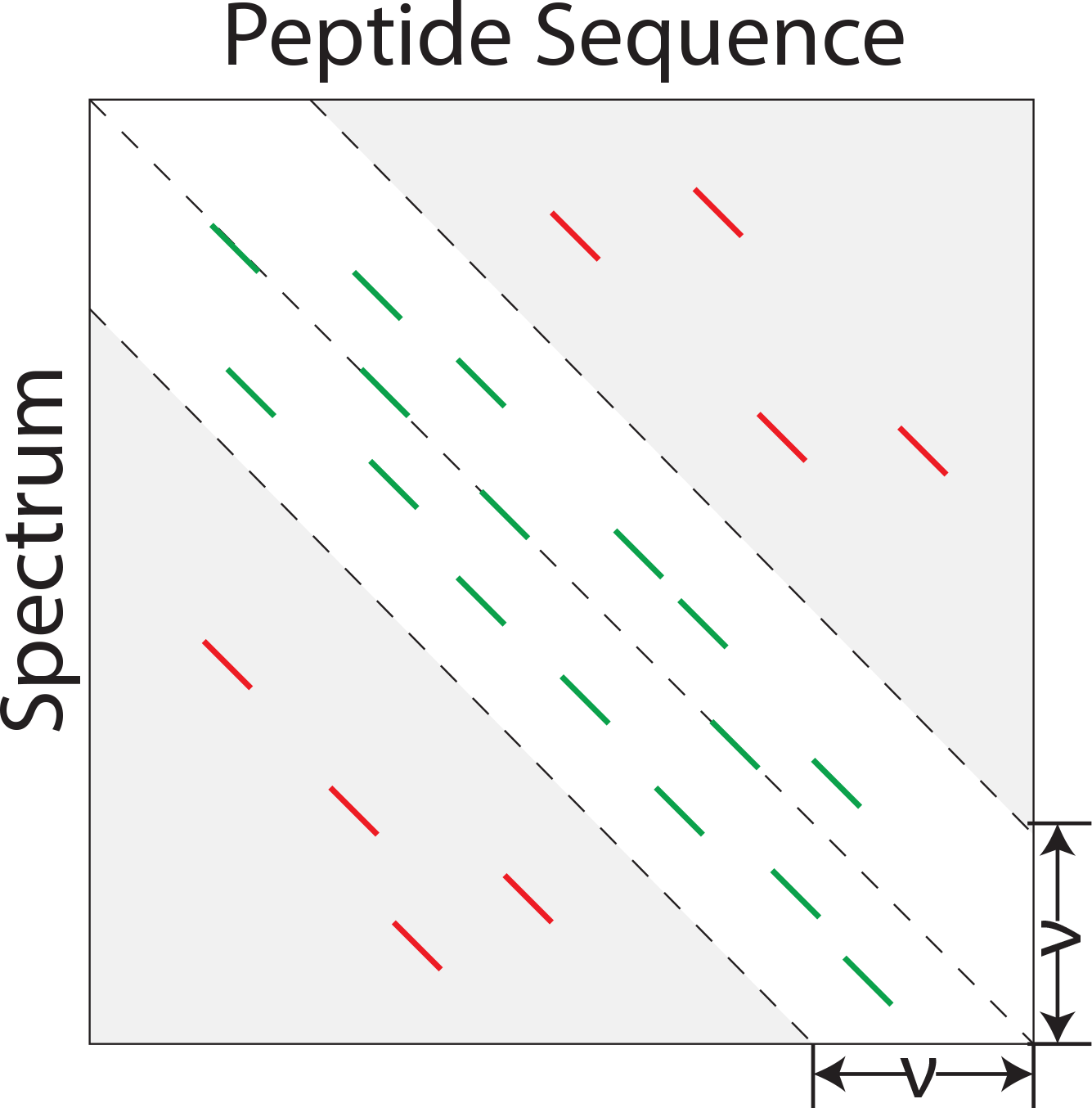
An illustration of tag cleaning. PIPI compares tags against a peptide sequence to obtain the relative shifts from the corresponding PTM-free location. The diagonal green or red bars indicate tags. PIPI only keeps those tags whose relative shifts are within a pre-defined range (green bars in the figure).

Jumps between tags can be classified into two categories:

1. There is no modified amino acid between two tags, which is called a non-PTM jump (circled 1 in Figure 6).
2. There are modified amino acids between two tags, which is called a PTM jump (circled 2 in Figure 6).

**Figure 6:**
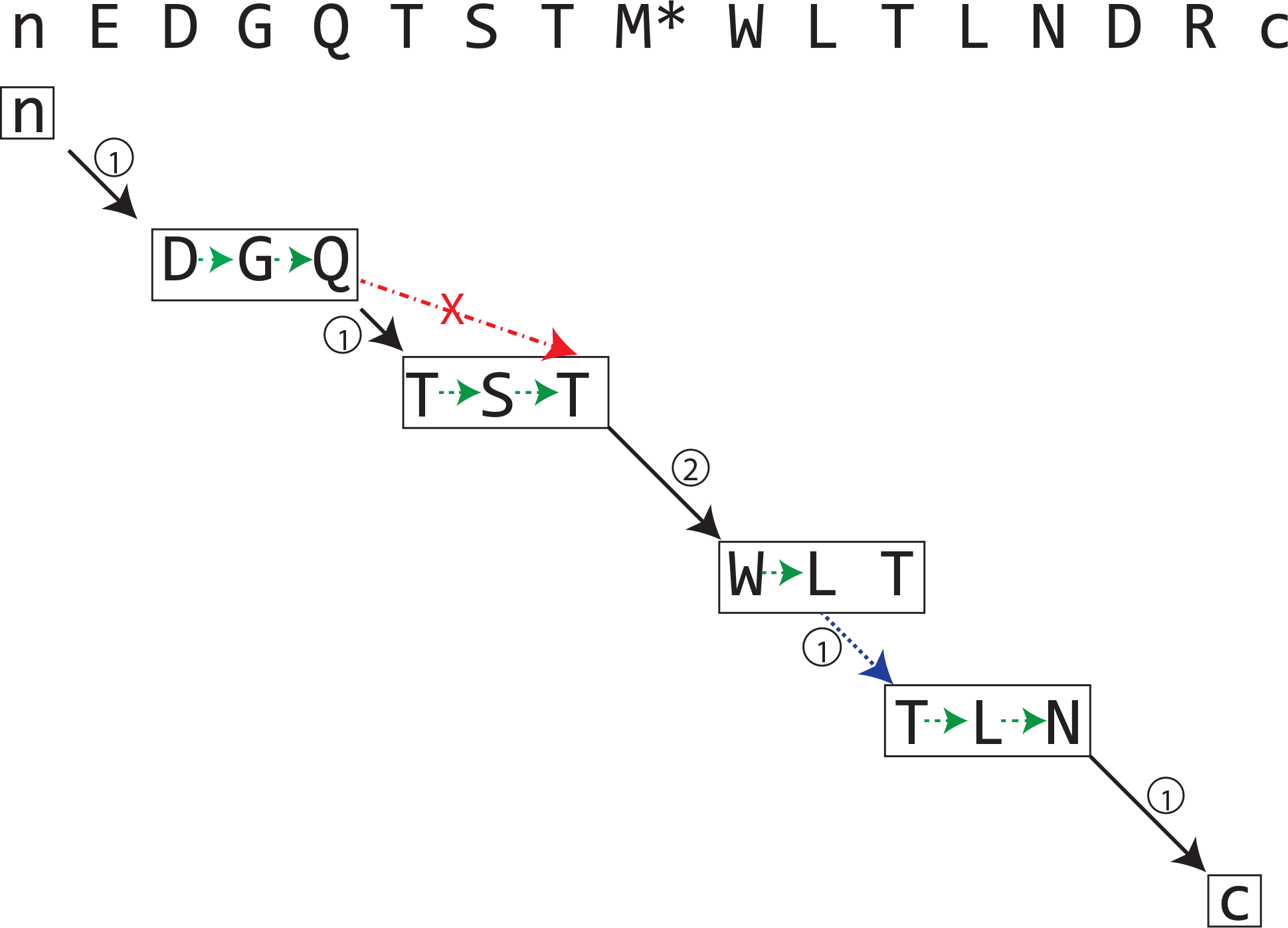
An illustration of tag alignment. Tags are aligned against a peptide sequence. The “n” and “c” indicate two special tags: X-terminal and C-terminal. There is a modification on “M” in the peptide sequence. Jumps between or within tags are labeled with different colors corresponding to different jumping types. Numbers on the jumping arrows indicate whether the jump is a non-PTM jump (circled 1) or a PTM jump (circled 2).

Thus, the scoring rules are as follows:

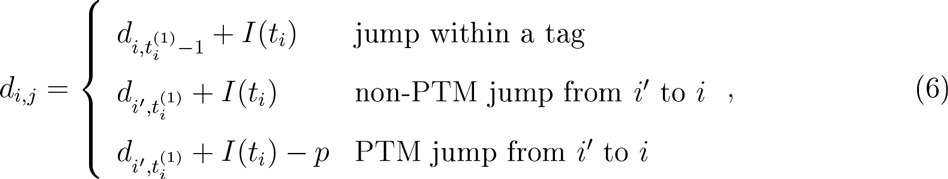
where *d_*i*,*j*_* is an element of **D**_|{*t_i_*}|×(*n*+2)_ and *p* is a penalty for a PTM jump.

### 2.4 Final scoring and *q*-value estimation

For peptide identification, restricted tools (e.g., Mascot, SEQUEST, MS-GF+, and Comet) have a higher sensitivity than unrestricted tools (e.g., MS-Alignment, ProteinProspector, and MODa) if modification patterns are included in the theoretical spectra. Besides, the original spectrum contains more information than the coded vector. As PIPI has already known each spectrum-peptide pair’s modification pattern after the PTM localization and characterization step (Section 2.3), it calculates scores using the original spectrum.

After the database search step (Section 2.2), each spectrum has 10 peptide sequences as candidates. PIPI calculates a score for each spectrum-peptide pair using the XCorr^2^:

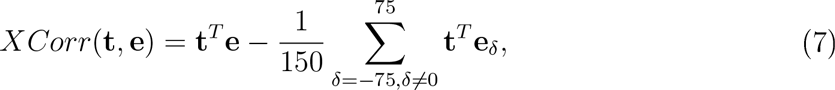
where **t** is a vector of the digitized theoretical spectrum, e is a vector of the digitized experimental spectrum, and *δ* is an *m*/*z* shift. XCorr is a score function used by popular tools such as SEQUEST and Comet. For each spectrum, the top-scored peptide is kept as the final result. Finally, we use Percolator^46^ to calculate the PSMs’ *q*-values. In most cases^46,49^, people convert FDR to q-value that is monotonically decreasing with respect to the score. Without specific description, we always convert FDR to *q*-value and use it as cut-off.

## 3 Experimental results

We used three sets of experiments to demonstrate the correctness and performance of PIPI. The first set contained 2 simulation experiments using 12,064 high-quality MS/MS spectra and two custom databases. The second used 5 public data sets from standard protein mixture samples^50^. The third used 24 public data sets from Chick et al. ^34^. Please refer to Klimek et al.^50^ and Chick et al.^34^ for details of the sample preparation and data acquisition.

In these three sets of experiments, we used MS-Alignment (version: 20120109), ProteinProspector (version: 5.16.0), MODa (version: 1.51), and PIPI (version: 20160418) to do the unrestricted search. MS-Alignment needs the maximum number of modifiable amino acids in each spectrum to be specified, and the default value is 1. We set it to 2 in the second simulation experiment and 1 in the other experiments. ProteinProspector works in either a restricted or unrestricted manner, and we used it in the unrestricted manner by allowing mass modifications on all amino acids. We set the maximum number of modifiable amino acids to 2 (the default value) in all experiments. MODa and PIPI don’t limit the number of modifiable amino acids in each spectrum. All of these four tools’ precursor mass tolerance was 10 ppm, and MS/MS mass tolerance was 0.02 Da. We only considered MS/MS spectra whose precursor masses were from 600 Da to 5000 Da. This is a common range, recommended by many tools^12,4,6,7^. The allowed modification delta mass was from −250 Da to 250 Da as in Chick et al.^34^. We allowed all amino acids, the N-terminal, and the C-terminal to be modified. We allowed no missed-cleavage. Because ProteinProspector doesn’t provide *q*-values for its results, we used an in-house program to estimate the *q*-values with the target-decoy strategy^51^. MS-Alignment and MODa do provide *q*-values for their results, and we used Percolator^46^ to estimate *q*-values for PIPI’s results. All four tools’ q-value cut-off was 0.01.

### 3.1 Simulation experiments

We picked 12,064 PTM-free MS/MS spectra from the data sets in Chick et al. ^34^. All of them have *E*-values≥0.01 as reported by Comet. The reason for using the *E*-value rather than *q*-value is that the *E*-value is more conservative and we would like to get highly confident results. These spectra correspond to 6,753 non-redundant peptides.

We randomly selected half of these peptides and randomly replaced one amino acid in each selected peptide according to the following rules:

1. “K” and “R” cannot be replaced.
2. “P” following “K” or “R” cannot be replaced.
3. Replaced amino acid cannot be “K” or “R”.
4. Replaced amino acid cannot be “P” if there is a “K” or “R” before it.
5. “I” cannot be replaced with “L” and vice versa.

With the modified peptides as a database, we had 6,111 spectra containing no modified amino acid and 5,953 spectra containing one modified amino acid. Let’s call this simulation data set “Simulation 1”.

We randomly selected half of the original peptides again and replaced two amino acids in each selected peptide at random. Then, we had another set of data containing 6,113 spectra without any modified amino acid and 5,951 spectra with two modified amino acids. Let’s call this simulation data set “Simulation 2”. The spectra files and databases can be downloaded from http://bioinformatics.ust.hk/pipi.html.

We added 116 common contaminant proteins from the common repository of adventitious proteins (cRAP)^52^ to the two databases, respectively. We also generated a decoy database by reversing the peptide sequences without changing the C-terminal.

Since we knew the ground truth of the two data sets, we could label the true positives and false positives for the results. Because ProteinProspector often reports multiple modification patterns for a PSM, we did not consider the difference in the modification patterns in the PSM comparison. Figure 7 shows the stacked bars of the results. For each bar, the yellow part corresponds to false positives and the blue part corresponds to true positives. The value in each blue part is the number of true positives, and the value at the top of each stacked bar is the total number of positives. PIPI identified more true PSMs than MS-Alignment, ProteinProspector, and MODa. We also calculated the false discovery proportion (FDP) for these results:

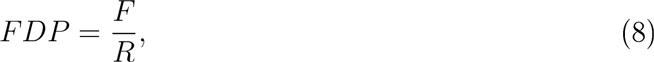
where *F* is the number of false positives and *R* is the number of positives. Table 2 shows the FDP of the results. PIPI outperformed the other three tools by providing more positive identification results with lower FDP values. The detailed results of these two experiments are available at http://bioinformatics.ust.hk/pipi.html.

**Table 2.** FDP of two simulation experiments. PIPI has the lowest FDP values.

| Tools \ Simulations | Simulation 1 | Simulation 2 |
| --- | --- | --- |
| MS-Alignment | 0.03 | 0.05 |
| ProteinProspector | 0.07 | 0.23 |
| MODa | 0.05 | 0.07 |
| PIPI | **0.02** | **0.03** |

**Figure 7:**
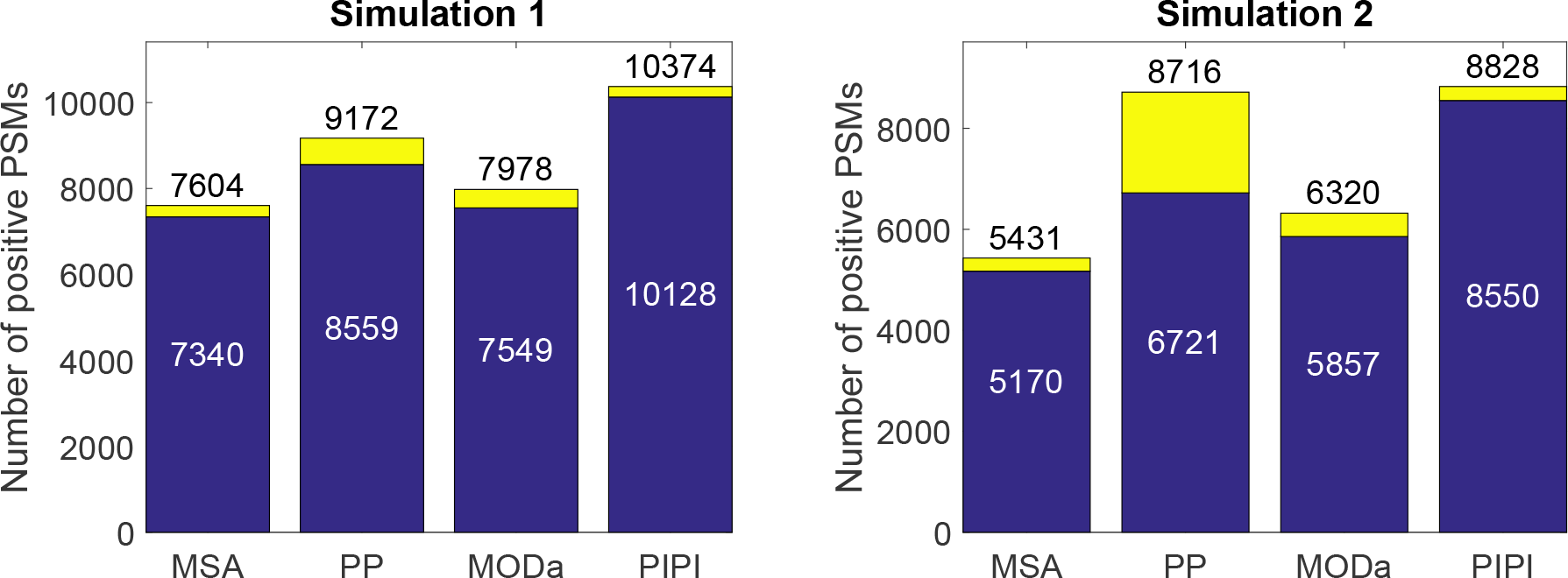
Bar plots showing the number of positive PSMs identified by MS-Alignment. ProteinProspector. MODa. and PIPI. respectively. “MSA” stands for MS-Alignment and “PP” stands for ProteinProspector. The first plot shows the results of “Simulation 1”. and the second plot shows the results of “Simulation 2”. The blue part corresponds to true positives, and the yellow part corresponds to false positives. The value in each blue part is the number of true positives, and the value at the top of each stacked bar is the total number of corresponding positives.

### 3.2 Experiments with standard protein mixture samples

We used five public data sets from the standard protein mixture samples ^50^ to demonstrate the performance of PIPI with real data. We used the database published along with the data sets, in which there are 18 standard proteins and 1.818 contaminant proteins.

Since the samples only contained 18 purified proteins, peptides belonging to these proteins had a highly possibility of being true positives, and peptides belonging to the contaminant proteins had a highly possibility of being false positives. We have plotted stacked bars showing the number of positive PSMs, as shown in Figure 8. For each bar, the yellow part corresponds to false positives and the blue part corresponds to true positives. The value in each blue part is the number of true positives and the value at the top of each stacked bar is the total number of positives. Since the decision on true positives was not accurate, we did not calculate the FDP for these results. These experiments showed that PIPI outperforms the other three tools in real data applications. The detailed results are available at http://bioinformatics.ust.hk/pipi.html.

### 3.3 Experiments with 24 real data sets

We used 24 data sets from Chick et al. ^34^. There are in total 1,309,561 MS/MS spectra whose precursor charges are from 1 to 7 and precursor masses are from 600 Da to 5000 Da. Since the samples were from HEK293 cells, we used the whole proteome of Homo sapiens from UniProtKB/Swiss-Prot (20,205 proteins, 2015–11 release) as the database for MODa and PIPI. MS-Alignment and ProteinProspector would need years to search these data sets against the whole human proteome, so we generated a small database based on the procedure proposed by MS-Alignment^17^. We first searched these data sets against the whole human proteome using Inspect^11^, which is a restricted tool. Then, we picked all the proteins that had at least 2 peptides and 10 spectra that were identified. We used these proteins to generate a small database, which, without considering decoy proteins, contains 4125 proteins. This approach was recommended by the authors of MS-Alignment and ProteinProspector.

Since we did not have the ground truth for the data sets, we only compared the number of positive PSMs. Because Chick et al. ^34^ only reported nonredundant peptides instead of PSMs, we did not compare our results with theirs. Figure 9 shows that PIPI identified more PSMs than MS-Alignment, ProteinProspector, and MODa in all data sets. This result is consistent with that from the last section. The detailed results can be downloaded from http://bioinformatics.ust.hk/pipi.html.

**Figure 8:**
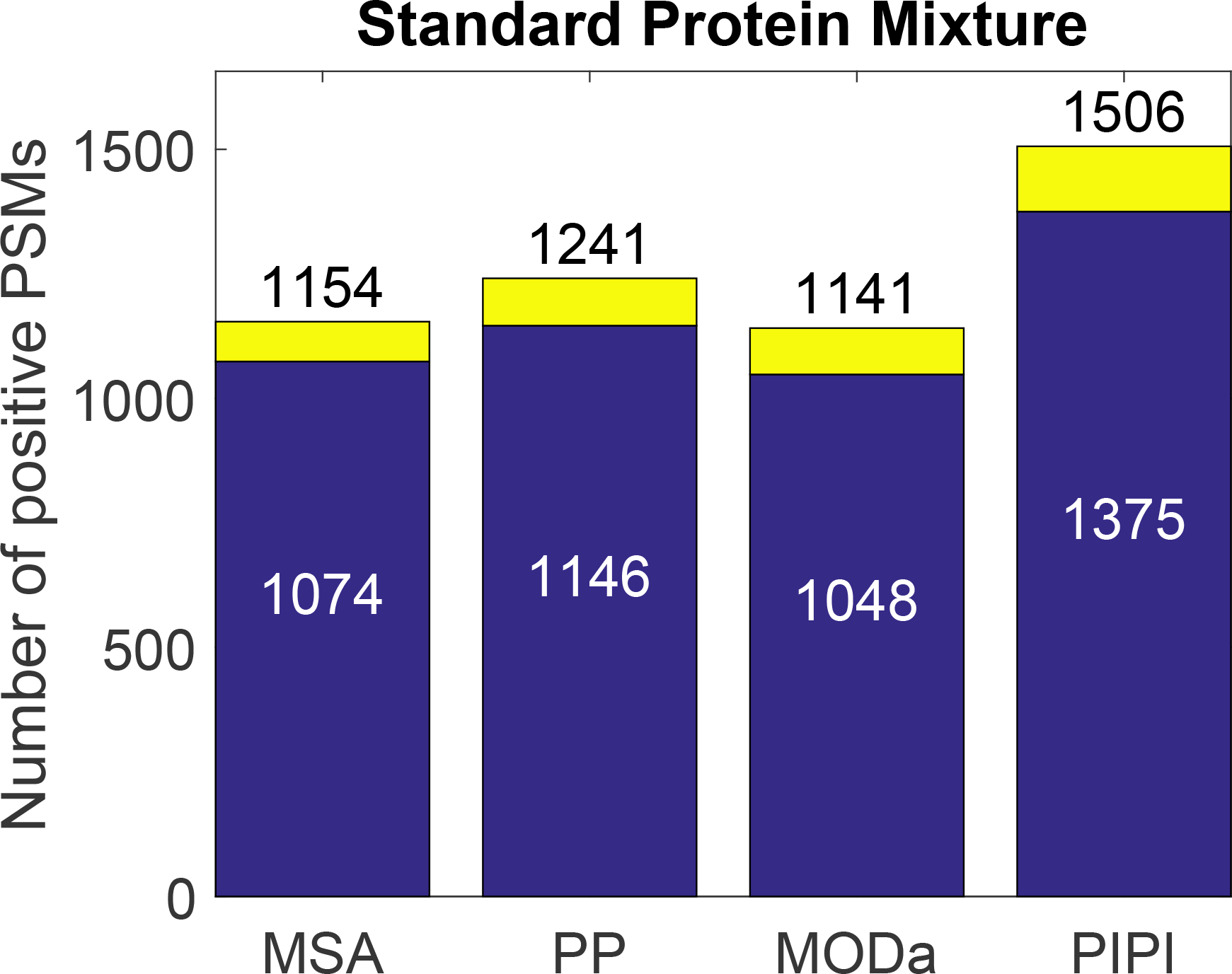
A bar plot showing the number of positive PSMs from identifying standard protein mixture samples. “MSA” stands for MS-Alignment and “PP” stands for ProteinProspector. The blue part corresponds to true positives, and the yellow part corresponds to false positives. The value in the blue part is the number of true positives, and the value at the top of the bar is the total number of positives.

### 3.4 Running time

We ran MS-Alignment. MODa. and PIPI on our computers with i7-6700 CPU (3.40 GHz) and 32 GB R AM. We ran ProteinProspector on the web server provided by its developers ^53^. As discussed in Section 3.3, we let MS-Alignment and ProteinProspector search against a small database, while MODa and PIPI searched against the whole human proteome. In the experiments discussed in other sections. MS-Alignment. ProteinProspector. MODa. and PIPI used the same database.

**Figure 9:**
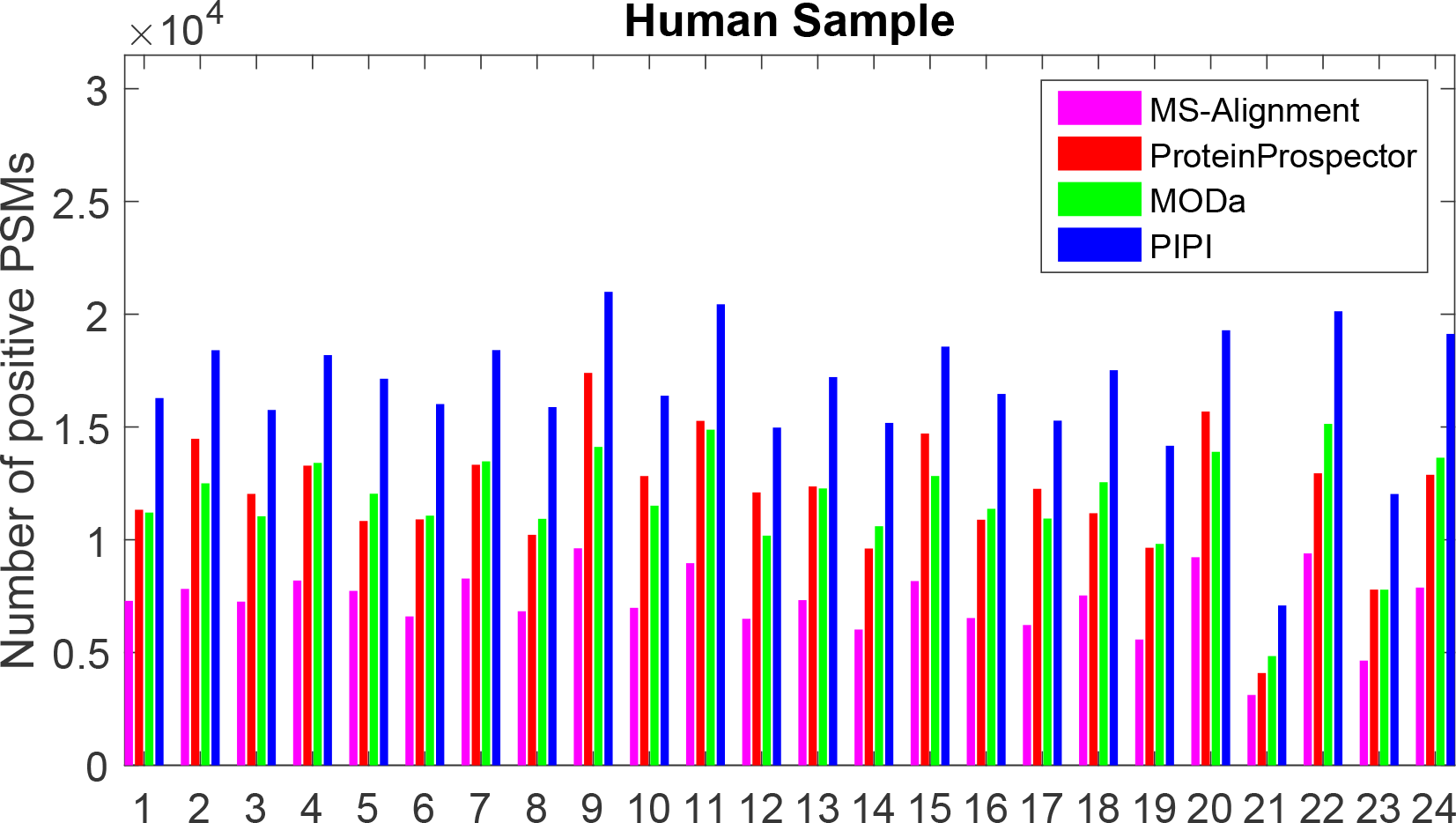
A bar plot shows the number of positive PSMs identified from 24 data sets. MS-Alignment. ProteinProspector. MODa. and PIPI were used.

**Table 3.** The average running time of analyzing one data set in the three sets of experiments, respectively. The unit is hours. The running time of MS-Alignment and ProteinProspector is marked with an “*” for analyzing 24 real data sets. This indicates that they used a custom database that is much smaller than the database used in MODa and PIPI.

| Tools<br>Experiments | MS-Alignment | ProteinProspector | MODa | PIPI |
| --- | --- | --- | --- | --- |
| Simulation Experiments | 15.05 | 0.77 | **0.07** | 0.25 |
| Standard Protein Mixture Experiments | 0.31 | 0.22 | **0.02** | 0.03 |
| 24 Real Data Experiments | 56.87* | 13.87* | 15.40 | **3.17** |

Table 3 shows the running time of analyzing one data set by MS-Alignment. ProteinProspector. MODa. and PIPI. respectively. The databases in the simulation experiments and standard protein mixture experiments were relatively small, while the databases in the 24 real data experiments were large. Table 3 indicates that PIPI is faster than all other tools in searching a large database.

## 4 Discussion

We can classify peptide identification methods into two main categories: *de novo* sequencing^36,38^ and database search ^13,6,7^. *De novo* sequencing infers a spectrum’s sequence without using any database. It checks pairs of peaks, and labels them if their distances are within the tolerance ranges of the amino acids’ masses. Then, it links the labeled peak pairs into paths and scores them. Finally, it finds a high-scored path and interprets the path into a peptide sequence. Clearly, *de novo* sequencing requires a high-quality spectrum. Missing peaks and unspecified PTM types are disasters for *de novo* sequencing. Database search infers a spectrum’s sequence by finding the most similar candidate from a database. After defining a scoring scheme, it compares each experimental spectrum with all possible theoretical spectra. The top-scored candidate is the final result. Clearly, this approach is tolerant to missing peaks and noisy peaks, but unspecified PTM types still cause trouble.

PIPI extracts local sequence information by inferring substring (a.k.a. tags) from a spectrum. The process of extracting tags is similar to *de novo* sequencing. But the key difference is that PIPI doesn’t try to infer the whole sequence. Instead, PIPI codes all tags into a feature vector and uses the feature vector for identification purposes. This procedure is similar to database search. The subtle difference is that database search compares an experimental spectrum with theoretical spectra, while PIPI compares a vector coded from a spectrum with vectors coded from peptide sequences. The former is sensitive to PTM, while the latter is invariant to PTM.

There are tools (e.g., MS-Alignment and MODa) that try to identify peptides without specifying PTM types beforehand. The major difference between these tools and PIPI is that the former perform alignment during the database search, while PIPI performs alignment after the database search. During the database search, MS-Alignment aligns an experimental spectrum against every possible theoretical spectrum, and MODa aligns a spectrum’s tags against every possible peptide sequence. These two tools use their alignment results in their scoring procedures. In contrast, PIPI represents experimental spectra and peptide sequences with coded vectors, and uses them to find each spectrum’s top 10 peptide sequences. After narrowing down the candidates, PIPI aligns a spectrum’s tags against each peptide sequence, and calculates a final score for *q*-value estimation.

**Table 4.** A table showing the numbers of PSMs having correct modification patterns, the total numbers of correctly identified PTM-containing PSMs, and their ratios.

| Tools \ Simulations | Simulation 1 |  |  | Simulation 2 |  |  |
| --- | --- | --- | --- | --- | --- | --- |
|  | True | Total | Ratio | True | Total | Ratio |
| MS-Alignment | 1540 | 2105 | 0.73 | 34 | 992 | 0.03 |
| MODa | 2172 | 3327 | 0.65 | 423 | 1717 | 0.25 |
| PIPI | 2816 | 4178 | 0.67 | 753 | 2596 | 0.29 |

There are also differences in the dynamic programming algorithms among these three tools. MS-Alignment aligns an experimental spectrum against a theoretical spectrum, while PIPI aligns tags against a peptide sequence. Because there are many peaks in an experimental spectrum, MS-Alignment is more than 10 times slower than PIPI, as presented in Section 3.4. MODa aligns variant lengths of tags against a peptide sequence, while PIPI aligns three-length tags against a peptide sequence.

As we mentioned in Section 1, the accuracy of PTM localization is low. We used two simulation experiments, as discussed in Section 3.1, to demonstrate this issue. Table 4 shows the numbers of correct modification patterns, the numbers of correct PSMs containing PTM, and their ratios. Because ProteinProspector often outputs multiple modification patterns for a PSM, we do not list its results in this table. Among the PSMs containing correct modification patterns, a considerable percentage have incorrectly characterized modification patterns. PTM localization is still an open question ^42–44^. The discussion of this question is beyond the scope of this paper.

## Acknowledgement

This work is partially supported by theme-based project T12-402/13N from the Research Grant Council (EGC) of the Hong Kong S.A.E. government.

## Supporting information available

The source code, executable file, simulation data sets, protein databases, and detailed results are available at http://bioinformatics.ust.hk/pipi.html.

